# Dynamics and spatial genomics of the nascent transcriptome in single mESCs by intron seqFISH

**DOI:** 10.1101/339234

**Authors:** Sheel Shah, Yodai Takei, Wen Zhou, Eric Lubeck, Jina Yun, Noushin Koulena, Eric J. Liaw, Mina Amin, Long Cai

## Abstract

Recent single cell experiments have revealed significant heterogeneities at the levels of transcription, DNA methylation and chromosome organization in individual cells. However, existing method of profiling mRNAs effectively averages transcriptional dynamics over many hours due to hours-long life time of most mRNAs. To capture the instantaneous activity of the transcriptome that reflects the rapid regulatory changes in cells, we imaged up to 10,421 nascent transcription active sites (TAS) in single mouse embryonic stem cells using seqFISH followed by multiple rounds of single molecule FISH and immunofluorescence. We observed that nascent transcription active sites appear to be distributed on the surface of individual chromosome territories and are dispersed throughout the nucleus. In addition, there are significant variability in the number of active transcription sites in single cells, representing globally more active to quiescent states. These states interconverted on the time scale of 2 hours as determined by a single cell pulse-chase experiment. Thus, transcriptome level seqFISH experiments provide an unprecedented spatial and dynamic view of chromosome organization and global nascent transcription activity in single cells.

## Introduction

Recent single cell experiments in mouse embryonic stem cells (mESCs) revealed heterogeneity at the transcriptional and protein levels (Chambers et al., 2007; Hayashi et al., 2008; Toyooka et al., 2008; Kumar et al., 2014; Klein et al., 2015; Kolodziejczyk et al., 2015), chromosome structures (Stevens et al., 2017) as well as epigenetic states (Hayashi et al., 2008; Marks et al., 2012; Singer et al., 2014; Smallwood et al., 2015; Angermueller et al., 2016). However, because most mRNAs have half-life of several hours in mammalian cells, mRNA levels average expression dynamics over many hours and obscure potential rapid changes in global transcriptional activity.

We set out to profile the global instantaneous transcriptional activity and chromosome structure of individual mESCs cells by Fluorescence in situ hybridization (FISH). We reason that instantaneous transcriptional profiling could reveal phenomenon occurring on a faster time scale compared to the longer-lived mature mRNA. In addition, FISH approaches can directly image gene loci to reveal the spatial organization of chromosomes in single nuclei.

Pioneering work with single molecule FISH (smFISH) (Femino et al., 1998; Raj et al., 2006) observed that nascent mRNAs are produced in bursts at transcription active sites (TAS) in individual nuclei. In particular, these nascent sites of transcription near the genomic loci can be selectively labeled over mature transcripts, by targeting introns, which are co-transcriptionally processed out (Levesque and Raj, 2013). This intron chromosomal expression FISH (iceFISH) assay (Levesque and Raj, 2013) showed that at least 20 TAS from a single chromosome can be detected to measure their spatial position and expression levels in single human cells.

Here, we applied the sequential FISH (seqFISH) strategy (Lubeck et al., 2014) to extend the iceFISH approach to the transcriptome level. seqFISH has been used to image hundreds of mRNAs in tissues and revealed distinct spatial structures in the mouse brain (Shah et al., 2016). The multiplex capacity of seqFISH is only limited by the density of the RNAs in cells and optical diffraction limit. Because nascent TAS occur infrequently and appear only near their genomic position in the nuclei, it is possible to resolve nascent site of synthesis across the genome in the same nucleus without optical crowding.

Furthermore, each TAS often consists of multiple nascent transcripts, enhancing the signal detected.

Using intron seqFISH, we profiled the instantaneous transcriptional activity of up to 10,421 genes in single mESCs, along with mRNA, long noncoding (lnc)RNA seqFISH and immunofluorescence to detect the pluripotency markers, cell cycle markers and nuclear bodies in the same single cells. We observed that active transcription occurs at the surface of chromosome territories and are distributed throughout the nucleus. We also observed that there are significant variability in the global transcriptional activities in individual cells. We showed that global instantaneous transcription activities are fluctuating rapidly on a 2 hour time scale using a single cell pulse chase experiment. The imaging approach taken here provides new information on spatial organization of loci and gene expression dynamics in single cells.

## Results

### Intron seqFISH targets transcriptionally active loci in single cells

To multiplex thousands of TAS loci, we used sequential rounds of hybridization to generate a unique temporal barcode sequence on each TAS, which can then be decoded by aligning images from each round of hybridization (Figure 1A-1B). Specifically, we target the introns at the 5’ regions of genes by a set of primary probes, which contain an overhang sequence that can be hybridized by the readout oligos that are labeled with fluorophores (Figure 1C). The cells are imaged on a spinning disk confocal microscope with z-sections. Then the readout probes are removed by denaturation in 70% formamide, while the primary probes remain bound on the intronic RNA due to longer probe size and higher DNA-RNA affinity. A different set of readout probes is then hybridized to the primary probes that switches the dye labels on each of the loci. Repeating the imaging, stripping and rehybridization N times provides *F^n^* barcodes, where F is the number of distinct fluorophores. With one additional round for error correction (Shah et al., 2016), we can correct for loss of signal in any round of hybridization due to mis-hybridization (Figure 1A).

**Figure 1.**
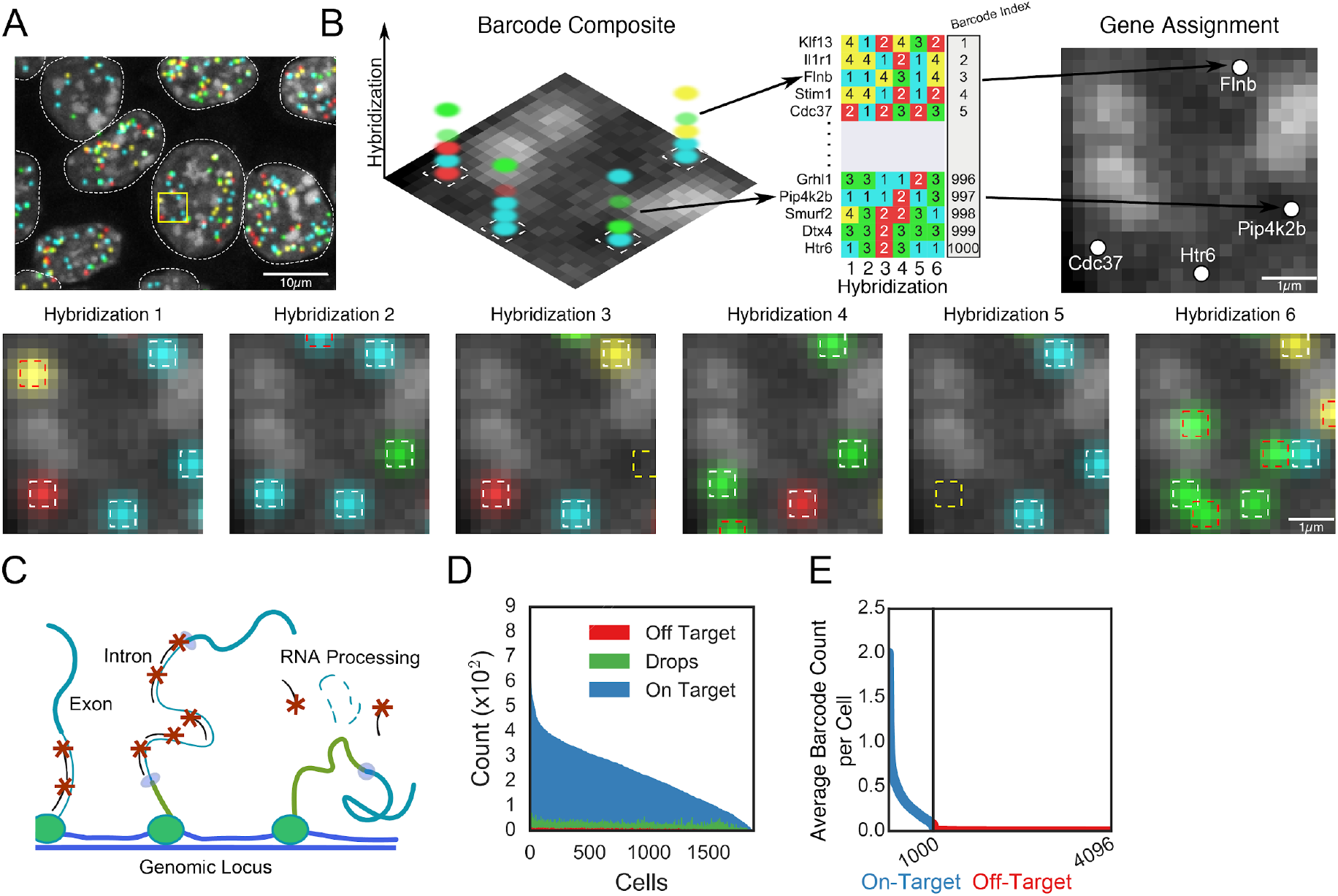
Intron seqFISH enables profiling of active transcription. (A) Large field of view image of multiple mESCs in a single hybridization from a single z-section. Below is a zoomed-in view of the boxed region over 6 sequential hybridizations. White boxes indicate identified barcodes, yellow boxes are recovered signal from error corrected barcodes, red boxes indicate false positive spurious hybridizations. (B) Identification of barcodes from zoomed region in A. The entire detected barcode is depicted as a concatenation of all 6 hybridizations. The genomic position of nascent RNA molecules is mapped by looking up barcode identities in a pre-assigned table. (C) Schematic depiction of intron FISH. Many nascent RNA molecules are present at the transcription active site, amplifying the detected signal. (D) Several hundred introns are typically detected per cell. Few off-target or incomplete barcodes are detected in any cell. (E) Average number of real (on-target) and false positives (off-target) introns detected per cell. Introns display a wide range of expression patterns, from near bi-allelic expression in every cell, to rare expression. False (off-target) introns are rarely detected, demonstrating the accuracy of intron seqFISH.

We first targeted the introns of 1,000 transcription factors that are conserved between mouse and human (Fulton et al., 2009) and are distributed across all of the chromosomes with 48 probes per gene. 6 rounds of hybridization and 4 fluorophores (4^6^=4096) can decode all 1,000 genes with one round of error correction. We observe negligible false positive rates in the decoding with the error correction scheme used (Figure 1B) with an average 188 ± 108 (mean ± standard deviation) on-target barcodes detected per celland 2.8 ± 2.3 (mean ± standard deviation) off-target barcodes detected per cell (Figure 1D). In individual cells, the number of active sites from each gene ranged from 0 to 4 per cell (Figure 1E), as some cells are in G2/M phase and contain 4 chromosomes. The estimated detection rate is 81% by comparing the number of dots detected in the first round of hybridization and the number of barcodes decoded. Compared to bulk GRO-seq data (Jonkers et al., 2014), which measures the amount of productively elongating RNA polymerase II and reflects the nascent transcriptional activity of genes, we observed a correlation of 0.57 with the average burst frequency of each gene in the seqFISH decoded data set. This comparison indicates the overall agreement between the burst frequency of active loci measured directly by intron seqFISH and the density of polymerases on gene loci measured by GRO-seq.

### Active transcription occurs on the surface of chromosomes territories

We next used the 1,000 gene intron seqFISH data to characterize the spatial organization of TAS in single mESCs. The data revealed that spatial distribution of TAS appears uniform across the nucleus, excluded from the DAPI dense heterochromatic regions as well as from the nucleoli (Figure 2A). There does not appear to be major factories of active transcription in the nucleus, although local foci cannot be ruled out. Reconstruction of the intron seqFISH data revealed that TAS from each chromosome appeared to span spatial territories (Figure 2A, right panel), consistent with previous observations (Mahy et al., 2002a; Bolzer et al., 2005) of chromosome territories (CTs) by chromosome paint experiments.

**Figure 2.**
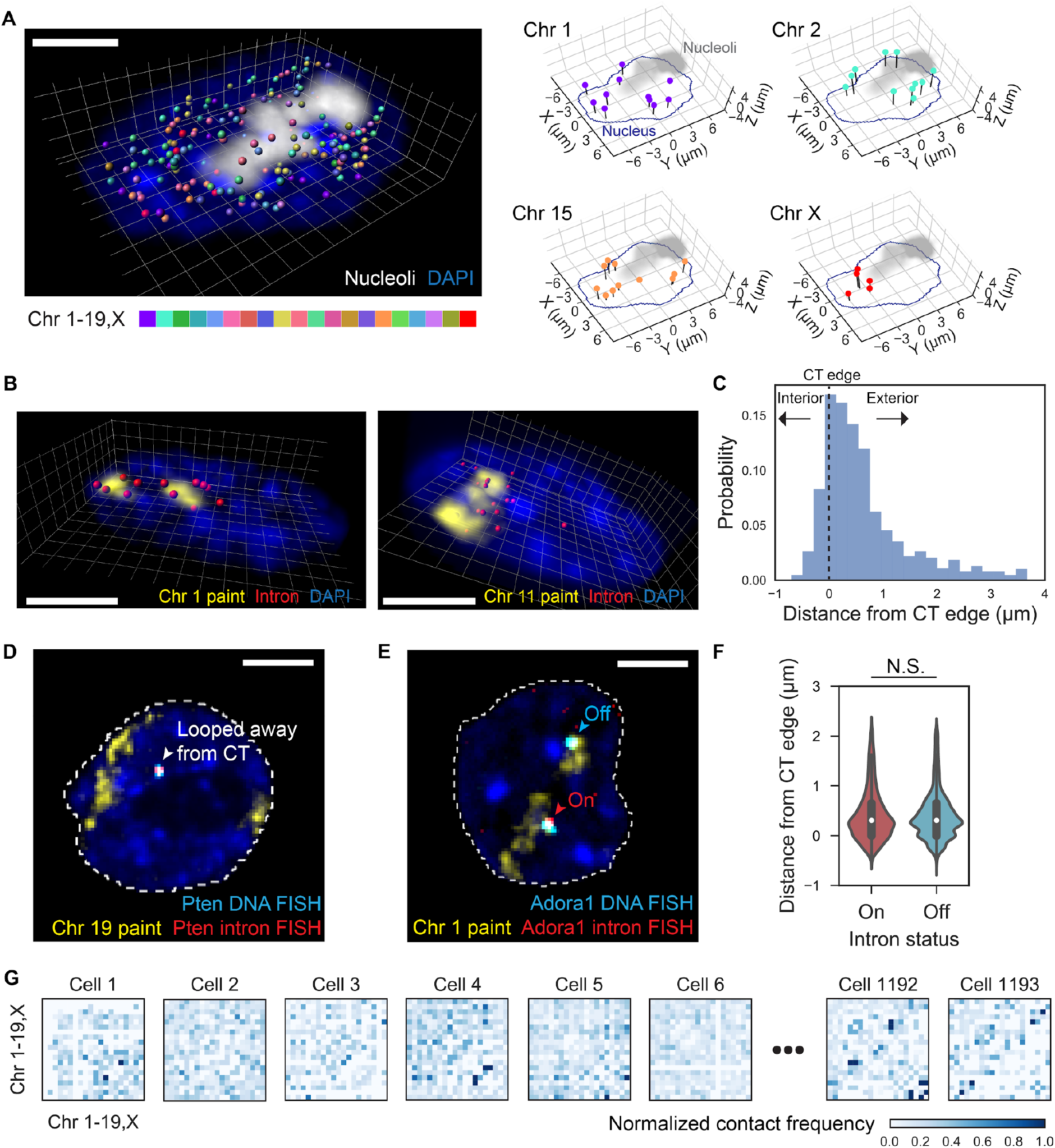
The 1,000 intron seqFISH reveals nascent transcription active sites at the surface of chromosome territories. (A) 3D reconstruction of TAS in a single mESC nucleus, with individual chromosome occupying distinct spatial territories (right). In total, 180 nascent sites were present in this cell. Nucleoli were labeled by ITS1 FISH and nucleus was stained by DAPI. (B) 3D reconstruction of introns from a particular chromosome (red) and chromosome paint of the same chromosome (yellow) in mESC nuclei (blue), showing nascent active sites present on the surfaces of chromosome territories. Chromosome 1 (left) and chromosome 11 (right) (intron probes targeting 77 and 79 genes on their chromosomes) are shown respectively. (C) Histogram showing the distance distribution of TAS relative to their chromosomal territory (CT) edge in mESC nuclei (n = 1472 spots from 4 chromosomes in total). (D) Locus looped away from its core CT, shown by DNA FISH (cyan) and intron FISH (red) targeting Pten along with chromosome 19 paint (yellow) in a mESC nucleus (blue). (E) Transcriptional status of loci does not affect their spatial positioning with respect to the CT boundary. DNA FISH (cyan) and intron FISH (red) targeted both allele of a gene (Adora1, as an example) along with chromosome 1 paint (yellow) in a mESC nucleus (blue). Signals outside nuclei (dashed white lines) are not shown for visual clarity (D, E). (F) Violin plots showing the distance distribution relative to its CT edge for loci with either “on” or “off” intron signals. N.S., not significant with Wilcoxon’s rank sum test (P > 0.05). Results from 9 genes in chromosome 1, including Adora1, are shown (n = 1550 and 4298 spots for “on” and “off” status). Scale bars (A, B, D, E), 5 μm. (G) Heat maps representing normalized contact frequency of TAS between pairs of chromosomes in single nuclei.

How are the TAS organized with respect to the CTs? To determine the relative positioning of the TAS and the CTs, we combined intron FISH targeting the TAS and chromosome paint probe hybridization to directly visualize the individual CTs. We observed that TAS are located on the surfaces of CTs (Figures 2B-D). We also found that occasionally some TAS can be located a significant distance away from the core CTs (Figure 2C-D). On average, TAS are located 0.65 ± 0.82 μm (mean ± standard deviation) exterior relative to their CT edge (Figure 2C). This is consistent with previous observations that regions containing coding sequences are positioned away from rest of the chromosome (Mahy et al., 2002b; Boyle et al., 2011). We found this feature is consistently observed across all profiled chromosomes, suggesting that surface RNA transcription is a universal feature of mESC chromosome. Several genes, such as Pten, Ppp3r1 and Ctnna1, showed lower intrachromosomal contacts compared to other genes, suggesting they are looped away from the main chromosome territory (Figure 2D). Individual intron FISH against these genes confirmed that they frequently loop out from their chromosomes and appear either on the surfaces of nucleoli or along the nuclear periphery (Figure 2D).

Next, we asked whether genes are dynamically localized to the surface of CTs depending on their activity. By performing intron FISH, DNA FISH and chromosome paint in sequential steps in the same cell (Figure 2E), we measured the spatial positioning of loci relative to the CT surface, as well as their instantaneous transcriptional activities. Notably, for the 9 genes we investigated, spatial positionings of loci were not influenced by their instantaneous transcriptional activities (Figure 2F).

We also asked how the TAS from different sets of chromosomes interacted with one another. As most of the TAS are present at the surface of the CTs or looped away from their core CTs, we observed many interchromosomal contacts and fewer intrachromosomal contacts. These interchromosomal contacts are highly variable at the single cell level, showing no consistent patterns (Figure 2G). These results are consistent with recent single cell Hi-C measurements (Nagano et al., 2013; Stevens et al., 2017). We note that intron seqFISH provides complementary information about the spatial organization of the nucleus compared to ensemble Hi-C (Lieberman-Aiden et al., 2009; Dixon et al., 2012), which captures features, such as topological associated domains, that are highly consistent amongst cells. As TAS are randomly distributed at chromosomal interfaces, their spatial organizations are averaged out in ensemble experiments.

### Transcriptome level intron seqFISH, mRNA, lncRNA seqFISH and immunofluorescence in single cells

We next sought to characterize the instantaneous transcriptional activity further at the transcriptome level as well as its relationship with the transcriptional states of the cell and nuclear bodies. To extend intron seqFISH to the transcriptome level, we designed probes targeted 10,421 genes and decoded these genes with 8 rounds of hybridization (4^8^=65,536; Figure 3A-B), which includes one round of error correction. We followed the intron seqFISH experiment with several addition rounds of mRNA, lncRNA seqFISH and antibody staining to target pluripotency, differentiation and cell cycle markers in addition to nuclear bodies in the same single cells (Figure 3A-B). We observed that 51 ± 29% (mean ± standard deviation) of the dots detected in the first round of hybridization are decoded per cell, with the lower rate of detection due to the fewer number of probes (24 per gene), limited by cost (a total of 250,104 probes were synthesized for all 10,421 genes), as well as loss of signal and alignment over 8 rounds of hybridization. The false positive rate for the 10,421 intron seqFISH experiment is estimated at 2.1 ± 1.2% per cell. We observed uniform spatial distribution of TAS in mESC nuclei (Figure 3C), consistent with the 1,000 gene intron seqFISH results (Figure 2A).

**Figure 3.**
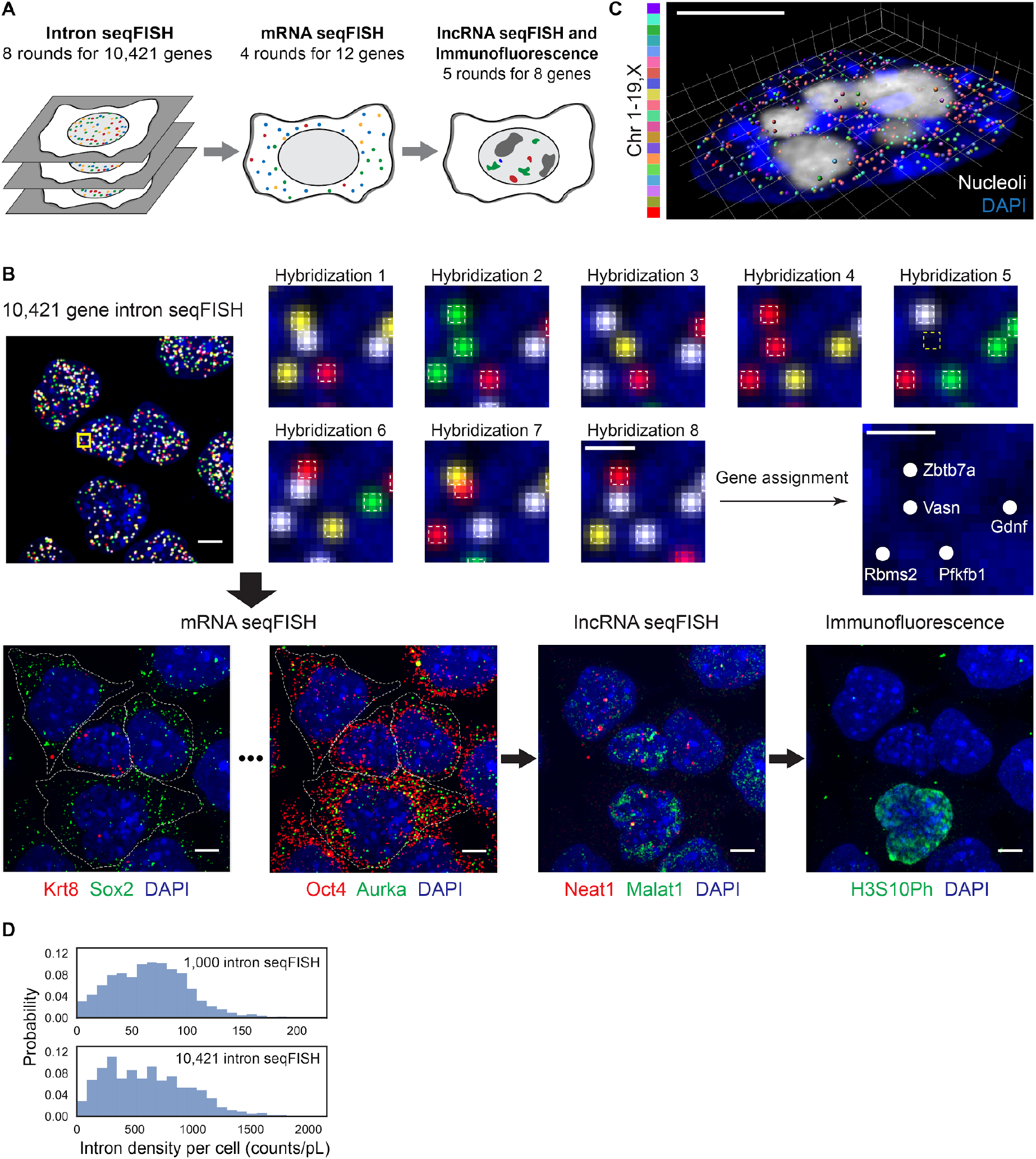
Transcriptome level seqFISH reveals heterogeneity in global nascent transcription. (A) Schematic of the combined seqFISH and immunofluorescence single cell measurements. 10,421 intron seqFISH decoded by 8 rounds of multiplexed barcoding were followed by additional rounds of mRNA, lncRNA seqFISH and immunofluorescence. (B) Combined seqFISH and immunofluorescence images with multiple mESCs. Barcodes in the boxed region of 10,421 gene intron seqFISH image are magnified. Colored boxes are depicted in the same way as Figure 1A. Note that intron barcodes are combined from 3 z-slices and digitized for visualization. Other images are maximum intensity projections of a z-stack of fluorescence images. Dashed white lines in mRNA seqFISH images display cytoplasmic boundaries of cells. Scale bars represent 5 μm in images with multiple single cells, and 1 μm in the magnified images. (C) 3D reconstruction of TAS from 20 chromosomes (chr1-19, X) in a single mESC nucleus. In total, 573 nascent sites were decoded in this cell by the 10,421 gene intron seqFISH. Nucleoli were labeled by ITS1 FISH and nucleus was stained by DAPI. Scale bar represents 5 μm. (D) Histograms showing wide distributions of nascent site density per cell from the 1000 gene intron seqFISH (top) and 10,421 gene intron seqFISH (bottom) (n = 1193 and 885 cells). Counts are normalized by nuclear volume (pL, picoliter).

Surprisingly, large variabilities in the overall instantaneous transcriptional states of cells are present, as determined by the number of active transcription sites in each nucleus in the 10,421 gene intron experiment (Figure 3D). There are on average 990 ± 634 TAS per cell (mean ± standard deviation), indicating that some cells are globally transcriptional active while other cells are globally quiescent. Similarly, wide distributions are also observed in the 1000 gene intron experiments (Figure 3D).

Notably, the variability in TAS number still exists within same cell cycle phases, which are distinguished based on immunofluorescence. Nuclear volume, on the other hand, is correlated with the total TAS numbers, but significant variations remain in each cell size window. Therefore, even when cell cycle and size variabilities are taken into account, there are substantial variabilities in global instantaneous transcriptional levels among single cells.

Using the lncRNA and immunofluorescence staining, we examined the spatial relationship between the TAS and nuclear bodies in mESCs. We observed that TAS are not strongly colocalized with nuclear bodies, such as the paraspeckle marked by Neat1, nuclear speckle marked by Malat1 and SC35, and lncRNA Firre, which controls a network of genes related to RNA processing in mESCs (Bergmann et al., 2015) (Figure 4A-B). Interestingly, while probes targeting polyA sequences are colocalized with SC35 speckles (Figure 4B and 4D), intron probes targeting TAS are weakly colocalized with SC35. We also observed heterogeneous expression of Malat1 and intermediate co-localization between Malat1 and SC35 labeled nuclear speckles. These observations are consistent with literature (Politz et al., 2006), and suggest that active transcription sites are not necessarily recruited to nuclear speckles.

**Figure 4.**
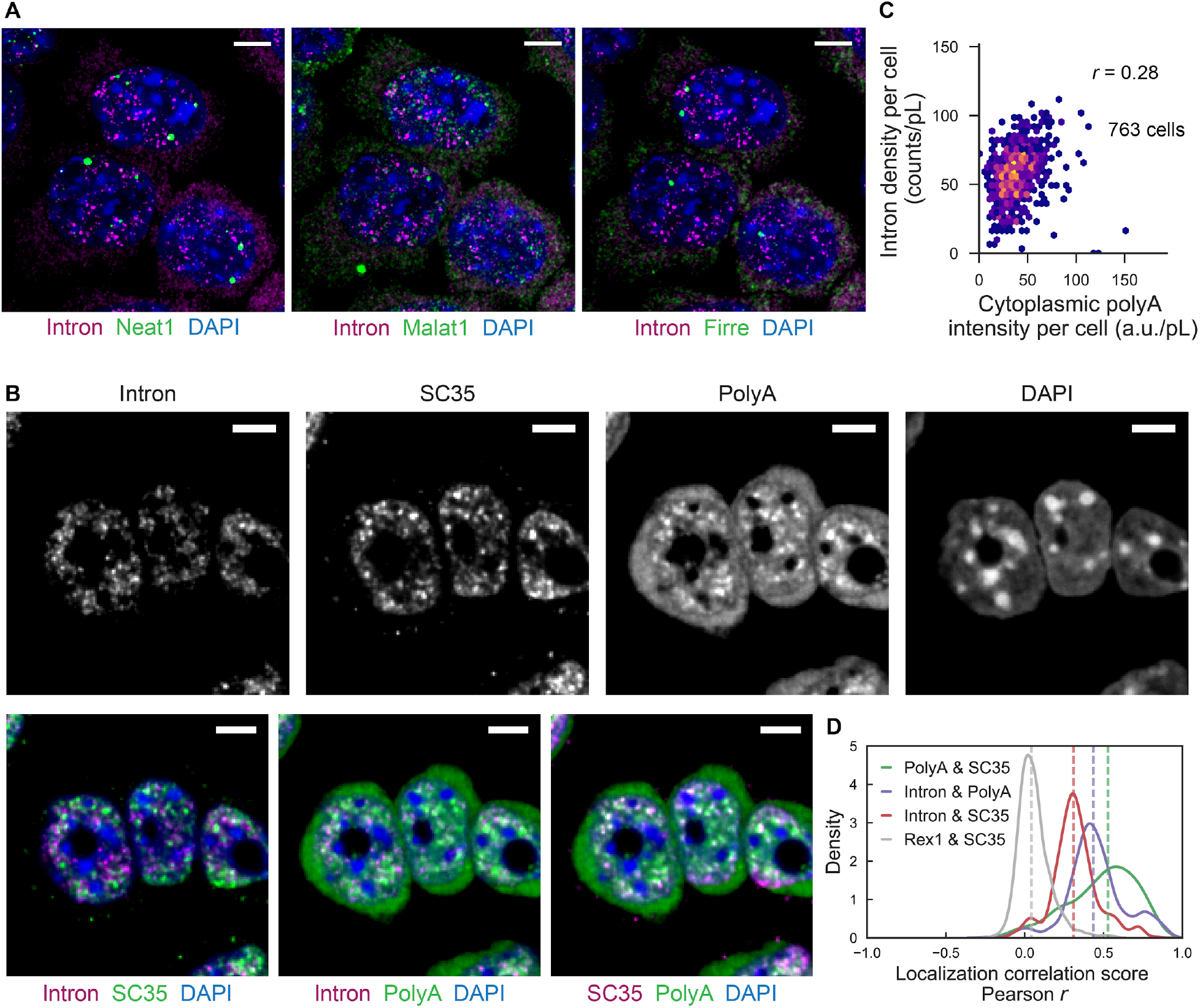
Organization of transcription active sites and nuclear bodies. (A) Representative images showing intron spots from the Alexa 488 channel in the first hybridization of the 10,421 gene intron seqFISH (magenta), lncRNAs by lncRNA seqFISH (green) and nuclear stain by DAPI (blue) in mESCs. Images are maximum intensity projections of a z-stack of fluorescence images. (B) Representative images showing intron spots from 1000 gene seqFISH, polyA FISH, SC35 immunofluorescence, and nuclear stain by DAPI. (C) Pairwise relationship between intron density by the 1000 gene intron seqFISH and cytoplasmic polyA intensity by polyA FISH (n = 763 cells). r, Pearson correlation coefficient. (D) Distributions of localization correlation scores (Pearson correlation coefficient) in single cells (n = 437 nuclei). Solid lines display density plots and dashed lines indicate median correlation scores from our data. Note that Rex1 (mRNA FISH) & SC35 correlation score represents baseline correlation. Scale bars (A, B), 5 μm.

### Global dynamics in transcriptional activities

The large variability in global transcriptional states raises the question of whether these global states are static in time or interconvert dynamically. Based on the weak correlation (R = 0.28) between the total TAS number in the nucleus and the amount of mature mRNA in the cytoplasm measured by dT oligos (Figure 4C), we hypothesized that the nascent global activities are not static in time. As the dynamics are unlikely to be synchronized amongst cells, we cannot measure the interconversion rate between active and inactive global transcriptional states by population averaged experiments. At the same time, it is also difficult to perform direct live cell experiments with reporter based assays to measure the transcriptional activities across many genes.

To overcome these limitations and measure the dynamics of TAS globally, we developed a single cell pulse chase experiment that records the nascent transcriptional activities at two time points in a cell’s history (Figure 5A). We first fed cells with a modified Uridine (5-EU) to record the global transcriptional activity during a short 30 minutes pulse. Then we washed out the 5-EU and let the cells grow for different amounts of time from 0 to 2 hours. We fixed the cells, measured the 5-EU incorporation levels with a clickable fluorescent dye and counted the number of TAS with the 1000 gene intron probes in the same cells. The variability in the 5-EU signal in individual cells is similar to the intron variability observed (Figure 3D), confirming nascent transcriptional heterogeneity in single cells.

**Figure 5.**
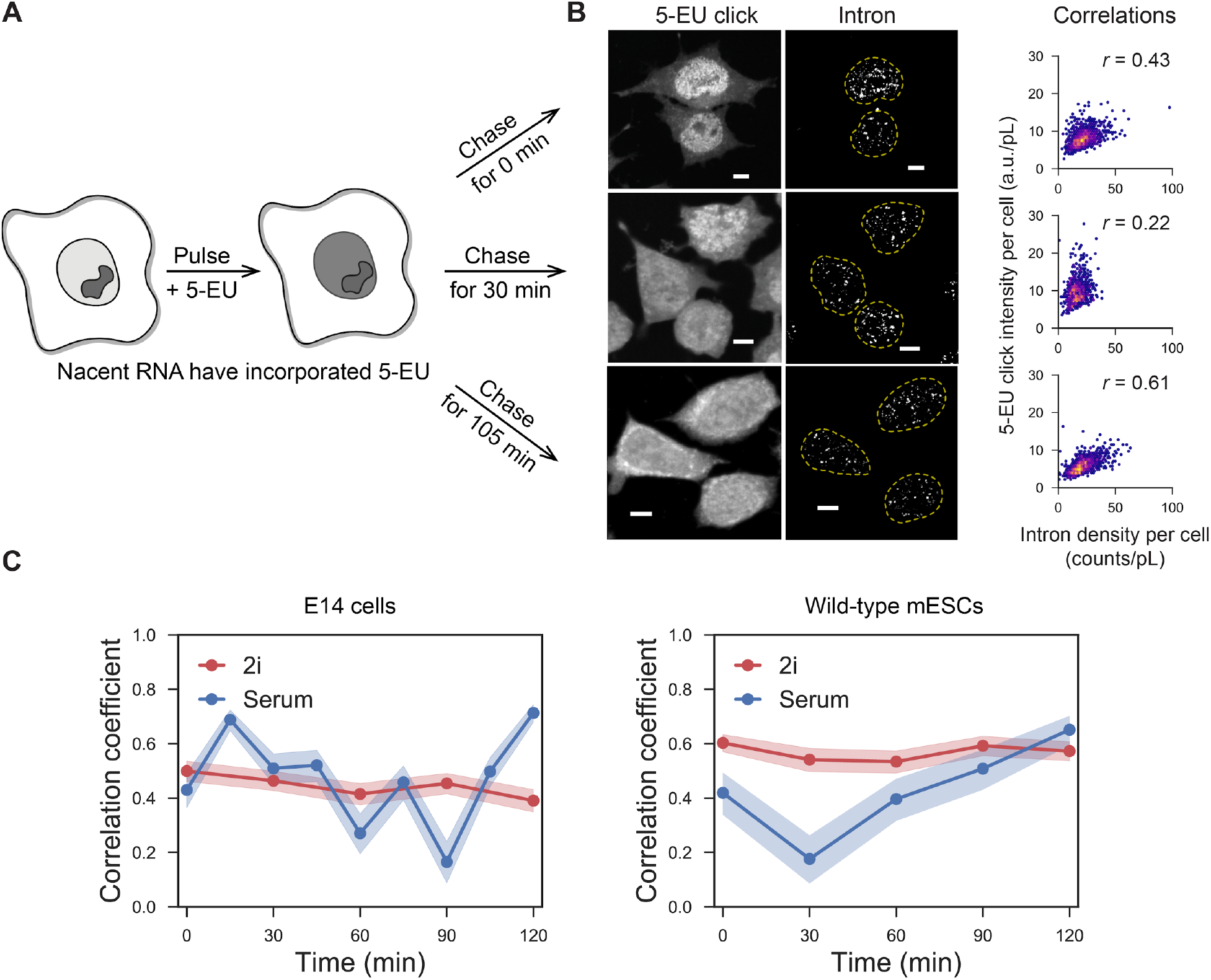
Single cell pulse chase experiments reveal fast dynamics in global nascent transcription. (A) Schematic of the pulse-chase experiment. 5-EU was pulsed for 30 min to globally label nascent transcripts, then chased for different periods of time in growth medium lacking 5-EU, followed by fixation. (B) Images of 5-EU detected by Click linkage to an azide-dye, and 1000 gene introns in single cells. The correlations of 5-EU and intron examined in different chase time were shown as scatterplots with Pearson correlation coefficient *(r)*, for 0 min, 30 min, and 105 min chase periods, respectively. (n = 1495, 780 and 1176 cells). Dashed yellow lines display nuclear boundaries determined by DAPI. Scale bars, 5 μm. (C) Pulse-chase correlation measurements show oscillatory dynamics on the time scale of two hours. Oscillations are observed in two cell lines of mESCs (left and right panels). In contrast to cells in serum conditions (blue), cells in 2i conditions do not show oscillations (red). Shaded regions represent 95% confidence intervals.

Next, we determined whether transcriptional activities are changing or static over time by comparing the global instantaneous transcriptional activity at defined time points in the past labeled by 5-EU incorporation with the activity at the present measured by intron levels in the same cells (Figure 5B). At early time points, the 5-EU and intron levels are correlated in single cells (Figure 5B, top panel), confirming that the heterogeneities observed in both measurements are consistent. The correlation coefficient decayed within 1 hours, with little correspondence between the 5-EU signal and intron levels in single cells (Figure 5B, middle panel). This indicates that the overall nascent activity in cells are highly dynamic, with the global activity decorrelating within a one hour time period. Surprisingly, the correlation is restored at around 2 hours (Figure 5B, lower panel, and 5C). This result suggests the transition of mESCs between low and high transcriptional states occurs with a 2 hour time period. Our data at each time point consist of hundreds of cells and the 2 hour oscillation is observed in multiple biological replicates as well as in a completely different line of mESCs (Figure 5C, right). It is also worth noting that the mean number of TAS per cell did not vary between pulse-chase time points, and only the correlations with the 5-EU levels at the single cell level oscillated.

This fast dynamics in the global nascent transcription is abolished in 2i condition (Figure 5C, red lines). It has previously been shown that mESCs in 2i conditions are globally hypomethylated and have homogeneous expression levels of pluripotency factors such as Rex1, Esrrb and Nanog (Singer et al., 2014). Our observations indicate that the mESCs in the naïve state in 2i conditions are static in time and that oscillation is an intrinsic feature of mESCs in the metastable pluripotent state in serum media where significant heterogeneity in transcriptional and epigenetic levels are observed (Marks et al., 2012; Singer et al., 2014; Kolodziejczyk et al., 2015).

Taken together, our finding of the global instantaneous transcriptional level fluctuations at the fast time scale is consistent with the lack of correlation with the mature mRNA levels in single cells (Figure 4C), as the longer lifetimes of mRNAs average out the fast dynamics in nascent transcriptional activities. As seen in a high degree of heterogeneity of the instantaneous transcriptional activities at any given time point, mESCs in the culture are not synchronized. However, cells interconvert between high and low instantaneous transcriptional states with a constant period of oscillation. If the global transcriptional activities were fluctuating stochastically without a defined period, or if different cells had different oscillation periods, then the correlation coefficients would simply decay without re-cohering at 2 hours.

## Discussion

In this study, we perform highly multiplexed intron seqFISH to capture the instantaneous activity of the transcriptome in large numbers of single mESCs. We combine these measurements with multiple rounds of mRNA seqFISH, lncRNA seqFISH and immunofluorescence to characterize transcriptional states, nuclear bodies and cell-cycle phases in the same single cells. This single-cell highly multiplexed imaging approach is complementary to sequencing-based methods such as GRO-seq (Core et al., 2008; Jonkers et al., 2014) and Hi-C (Lieberman-Aiden et al., 2009; Dixon et al., 2012) in measuring the nascent transcriptional activity and chromosome organization, respectively. Even though the seqFISH approach does not have nucleotide level resolution on chromosomal loci, imaging allows direct mapping of spatial organization beyond pairwise interactions measured in Hi-C and provide spatial context with respect to nuclear bodies through immunofluorescence and lncRNA staining in the same cells. In addition, the single molecule based seqFISH methods allows sensitive detection of nascent transcripts in single cells, currently not available with sequencing methods. Lastly, combining intron FISH with pulse labeling provides dynamic information that would be otherwise lost in population average measurements or in reporter-based single gene live cell experiments.

The spatial genomics approach provides a systematic view of the organization of nascent transcription active sites in mESCs (Figure 6A). Specifically, we found that first, TAS are in general distributed at the surface of chromosome territories, with some loci looped away from the CTs, and spread across the nucleus with the exclusion of nucleoli and heterochromatic regions. Second, TAS in different chromosomes can be located in close proximity and these inter-chromosomal contacts are highly variable among single cells, consistent with reports that regions where chromosomes intermingle are enriched with transcriptionally active phosphorylated RNAPII in mESCs (Maharana et al., 2016). Third, instantaneous transcriptional activity does not affect spatial positioning of the genomic loci relative to their CTs, and those loci appear at the surface of CTs even they are transcriptionally quiescent. This is consistent with previous reports that instantaneous transcriptional differences may not affect chromosome positioning from nuclear periphery (Levesque and Raj, 2013) and between homologous alleles (Takebayashi et al., 2012). However, it is possible that changes of transcriptional activity could drive reorganization of chromosome conformation at specific cases or regions such as active and inactive X chromosomes (Naughton et al., 2010), and HoxB cluster (Chambeyron and Bickmore, 2004). Further investigation on these specific loci can take advantage of intron seqFISH together with multiplexed live cell imaging of genomic loci (Takei et al., 2017).

**Figure 6.**
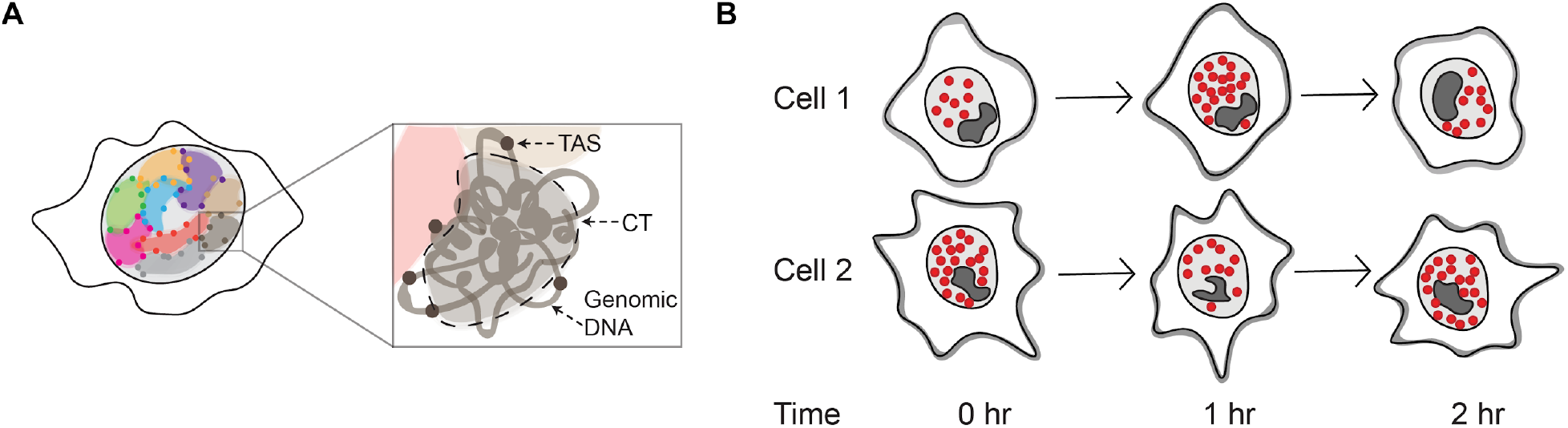
A model for spatial organization and dynamics of global nascent transcription in mESCs. (A) transcription active sites (TAS) are organized at the surface of chromosomal territories (CTs) and interfaces. (B) Oscillatory dynamics in global nascent transcription activity in two different cells. Red dots indicate TAS. Dark patches represent nucleoli.

Our data further showed that instantaneous global transcriptional activities have large variabilities amongst cells. These global states oscillate on the time scale of 2 hours in mESCs cultured under metastable serum + LIF conditions (Figure 6B) but not in naïve 2i conditions, where mESCs are known to be less heterogeneous and have reduced DNA methylation levels (Singer et al., 2014). Indeed, a recent study discovered that DNA methylation also oscillates with 2 hour periods in mESCs released from naïve into primed culture conditions (Reik, personal communication). Furthermore, it has reported that Hes1, Dll1, and Gadd45g promoter activities oscillated on a 2-5 hour time scale in mESCs (Kobayashi et al., 2009). These observations suggest that global oscillations in both DNA methylation and transcriptional activity may be connected and should be further explored.

The unexpected dynamics in the global transcription activities in mESCs shows that single cell whole transcriptome profiling and pulse chase experiments can reveal hitherto unknown regulatory dynamics. Using dynamic regulatory mechanisms, cells can achieve states not accessible with amplitude based regulation schemes (Letsou and Cai, 2016). For example, cells can use fluctuations in global transcriptional activity to coordinate the stoichiometry of many transcripts in a mechanisms akin to the frequency modulated signaling observed in yeast and mammalian pathways (Cai et al., 2008; Yissachar et al., 2013). It will be fascinating to determine the mechanisms underlying this oscillation and investigate whether similar fast dynamics occur in the inner cell mass of embryos.

